# Why be thrifty? Sex-specific heterothermic patterns in wintering captive *Microcebus murinus* do not translate into differences in energy balance

**DOI:** 10.1101/2021.08.21.457194

**Authors:** Aude Noiret, Caitlin Karanewsky, Fabienne Aujard, Jeremy Terrien

## Abstract

The physiological mechanisms of the responses toward stressors are the core of ecophysiology studies to understand the limits of an organism’s flexibility and better predict the impact of environmental degradation on natural populations. However, little information is available when we question inter-individual variability of these physiological responses, even though they can be particularly important. Some observations of intersexual differences in heterothermy raised the question of a difference in energy management between sexes. Here we assess male and female differences in a mouse lemur model (*Microcebus murinus*), a highly seasonal Malagasy primate, studying their physiological flexibility toward caloric restriction, and examining the impact on their reproductive success. These animals are adapted for naturally changing food availability and climate conditions, and can express deep torpor, allowing them to spare their energy expenses over the dry and cold season. We monitored body mass and body temperature on 12 males and 12 females over winter, applying a chronic 40% caloric restriction to 6 individuals of each group. Our results showed variability of Tb modulations throughout winter and in response to caloric treatment depending on the sex, as females entered deep torpor regardless of food restriction, while only CR males had a similar response. The use of deep torpor, however, did not translate into better body condition either in females, or in response to CR, and did not clearly affect reproductive success. The favorable captive context potentially buffered the depth of torpor and minimized the positive effects of using torpor on energy savings. However, our results may emphasize the existence of other benefits of heterothermic responses than fat reserves.

## Introduction

The physiological mechanisms of the response toward stressors are the core of ecophysiology studies to understand the limits of an organism’s flexibility in a context of environmental perturbations. Seasonality is a ubiquitous feature which has driven the evolution of many life history traits in the majority of species (Lisovski, Ramenofsky, and Wingfield 2017; Moen, Andersen, and Illius 2006; Williams et al. 2017). The succession of seasons implies that organisms are capable of preserving homeostasis in either periods of low or high resource availability, with strong variations in abiotic features such as ambient temperature or rainfall. Various responses, such as migration or hibernation, allow animals to adapt to circannual fluctuations, and all require metabolic flexibility (Canale and Henry 2010). Whether these responses induce high metabolic rate to trigger large-scale movement in order to find suitable energy resources elsewhere, or hypometabolism to save energy during low food availability, animals all engage in an energy compromise that maintains survival and ultimately sustains reproduction. Indeed, many species organize their reproductive schedule around the most favorable period to support juvenile growth and survival, and thus invest energy allocation in different biological tasks accordingly. To buffer the costs of an unfavorable season on fitness, many species store fat and share the strategy of “capital breeding” (Doughty and Shine 1997; Varpe et al. 2009). Still, because mating must be prepared for in advance in order to produce gametes, and is then followed by the development and growth of the offspring, energy expenses vary in amplitude while occurring in more or less challenging environmental conditions.

However, little information is available when we question inter-individual variability of energy balance strategies, even though they can be particularly important. Some observations of intersexual differences in torpor and hibernation abilities raised the question of an effect of sex in energy management. In the little brown bat (*Myotis lucifurus*), females manage to retain more fat going through winter than males, which led to the formulation of a “thrifty female hypothesis” (TFH), that would confer an advantage to females upon environmental challenges (Jonasson and Willis 2011; Czenze, Jonasson, and Willis 2017). In polar bears (*Ursus maritimus*), only females enter hibernation when they are in gestation (Lennox and Goodship 2008), and adult male ground squirrels express shorter torpor bouts than females (Kart Gür and Gür 2015). For long-day breeders, these sex-dependent expressions of hypometabolism are likely to be linked to males’ own agendas of spermatogenesis, which requires energy investment during winter before the mating season (Gagnon et al. 2020; Barnes et al. 1986). Females, on the other hand, are known to reactivate their reproductive axis by responding to photoperiodic change when transitioning to summer (Perret and Aujard 2001; Simonneaux and Ribelayga 2003). As a consequence, while males produce high quantities of gametes during low food availability, females stay reproductively and metabolically quiescent and spare energy stores until the favorable season. However, females’ allocation to reproduction is largely thought to be greater than males’, especially because of lactation in mammals, which is recognized as requiring the most energy expenditure (Canale, Perret, and Henry 2012; Clutton-Brock, Albon, and Guinness 1989). Nevertheless, although females will mostly rely on their fat reserves to fuel such demanding process, they can also compensate high energy expenditure with increased foraging in an energy-favourable context. All of these factors are likely to cause a gap of energy balance between sexes during winter, either from an adaptation to high reproductive expenses in females, or from a direct consequence of active metabolism during low food availability for males.

Evidence of another case of thrifty females has been shown in grey mouse lemurs (*Microcebus murinus*), a highly seasonal and heterothermic Malagasy primate able to enter deep torpor in winter (Génin and Perret 2003; Giroud et al. 2008). Interestingly, males and females express strong differences in their body mass fluctuations over winter, as males begin to lose weight around the middle of the season concomitantly to testicular growth initiation (Perret and Aujard 2001; Terrien 2018), while females keep their fat storage until summer, when mating takes place. In nature, females store fat before entering winter and can lose up to 31.7% of their body mass during the season. In contrast, males’ capacity to gain weight seems to be weaker and their body condition stays approximately the same between the beginning and the end of the “inactive” phase (Schmid 1999). Moreover, females were described as using deeper and longer torpor bouts than males to face food rarefaction at the onset of winter season in field conditions (Vuarin et al. 2015). The expression of sex-specific thrifty phenotypes during winter could thus very well depend on environmental conditions, as shown by the different responses of captive and natural populations. We already assessed major sex-differences in *M. murinus’* physiological response to CR during summer (Noiret et al. 2020), but no experimental study has yet directly compared sexes throughout winter, although they are expected to adapt their metabolism differently depending on their specific reproductive tasks. We monitored body mass and body temperature on 12 males and 12 females over the entire 6-month short-day period (SD), applying a chronic 40% caloric restriction to 6 individuals of each sex group. By doing so, we aimed to explore the link between food shortage and the expression of sex-specific hypometabolism during winter. Indeed, we expected to find optional use of torpor in late winter especially in males, constrained to elevate their metabolic rate to perform spermatogenesis before the photoperiodic transition to summer. As torpor has been demonstrated to have indirect costs (Landes et al. 2020), the use of deep or frequent torpor could be inefficient – in terms of energy savings – as already exemplified in sexually active male mouse lemurs (Villain et al. 2016). We thus established daily profiles of body temperature at 3 different time points of the winter follow-up (early, middle and end of winter), in order to characterize males’ and females’ particular metabolic fluctuations throughout winter. As we expected females to conduct deeper and longer torpor bouts than males, we hypothesised they would lose less body mass than males in response to CR, confirming the TFH. Because food shortage is a usual environmental challenge during winter, we did not expect any impact on reproductive success, either on males or females.

## 2. Materials and Methods

### Tested animals and ethical concerns

Twenty-four grey mouse lemurs (*M. murinus*), 12 males and 12 females, all aged from 2 to 4 years (2.75 ± 1.07 years) and raised in the breeding colony of Brunoy (MNHN, France, license approval n° E91-114-1), were included in the experiment. Animals were kept in individual cages of 66 × 50 × 60 cm, equipped with branches, foliage and wooden nesting box, in semi-isolation from the others (visual, auditory, and odorant interactions remaining possible between individuals) for the duration of the experiment. Temperature and humidity were maintained constant (24-26°C and 55% respectively). Photoperiod was set to mimic winter as light exposure was restrained between 7am and 5pm for the 6-month duration of the experiment (short days, SD; 10 hours of light, 14 hours of dark, for 27 weeks). The lemurs were fed with a standard fresh mixture (egg, concentrated milk, cereals, spice bread, cream cheese and water), and banana, and were provided with *ad libitum* water. All described experimental procedures were approved by the Animal Welfare board of the UMR 7179, the Cuvier Ethics Committee for the Care and Use of Experimental Animals of the Muséum National d’Histoire Naturelle and authorized by the Ministère de l’Enseignement Supérieur, de la Recherche et de l’Innovation (n°14075-2018031509218574) and complied with the European ethical regulations for the use of animals in biomedical research.

### Experimental design

Animals were fed every day of the week, except on Saturdays, when they were fed twice as much for the weekend (food was not provided on Sundays). They received a control ration (Control treatment ‘CTL’) of 22g of mixture and 7g of banana (28.21 kcal.day^-1^) for one month at the onset of winter. After the first month, animals were divided into two groups of 12 mouse lemurs (6 males, 6 females), which received two different caloric treatment (‘Trt’ effect). One group was kept under CTL feeding for the rest of the experiment, and the other was fed daily with a 40% reduced ration until the end of winter (Caloric Restriction ‘CR’, 16.92 kcal.day^-1^). They were monitored with once per week weighing to follow their body mass (‘BM’ in g) and evaluation of their general condition. To avoid extreme fattening patterns, all feeding regimes were reduced by 20% on the 12^th^ week of winter (CTL = 22.57 kcal.day^-1^; CR = 13.54 kcal.day^-1^). Body size (BS) was measured from the tip of the nose to the anus in order to calculate Body Mass Index (BMI= BM/BS, in g.cm^-1^), and testis size was monitored once a week for males (rated from 0 to 2 based on their size and consistency: 0= testes are up in the abdominal cavity and scrotal sacs are loose; 0.5= testes descend and measure 1 cm put together; 1= 2 cm; 1.5 = 3cm; 2= 4cm and consistency becomes harder).

### Body temperature

Mouse lemurs were implanted with ANIPILL capsules (Bodycap, Phymep) into their abdominal cavity. Surgical procedures were performed under general anesthesia (Valium 0.5mg/ 100g; Ketamine 2 mg/ 100g; isoflurane maintenance) and per-operating analgesia (before surgery: buprenorphine 0.05mg/kg IM 30 minutes, local cutaneous injection of lidocaine around the abdominal aperture; after surgery: renewal of buprenorphine 4 hours later, then meloxicam *per os* for 2 days). After 11 months of recording, the implants were easily removed by the same procedure and no inflammatory lesion was observed. This method allowed the recording of each individuals’ body temperature (Tb) every 15 minutes. Tb was analyzed over the entire winter with minimal Tb registered each day (‘Tmin’, in °C), and its time of expression (‘Hmin’, in minutes from light switch off at 5pm). We completed the study of body temperatures with a phenological analysis of torpor use, where each individual was considered successfully using deep torpor as long as its weekly averaged Tmin was below 33°C (Génin and Perret 2003).

To better describe the use of heterothermy in these animals, median Tb variations were also analyzed over a 24-hour period of time during early (week 2 to 3), mid-(week 9 to 14) and late (week 22 to 25) winter periods. We used the ‘Segmented’ 1.1-0 package (Muggeo 2019) to estimate 5 linear regressions (and 4 breakpoints) from the median Tb daily curves, starting at 8pm (3 hours after night initiation) to keep the entire heterothermic profile uncut. For each individual and each of the 3 periods, several parameters were extracted from the curves to implement “heterothermy profiles” (Figure 1): time of heterothermy initiation (‘Hi’, in minutes from 8 pm), as the estimate of the most negative slope before minimal Tb; minimal Tb (‘Tmin’, in °C) as the lowest of breakpoints ordinates, and its hour of expression (‘Hmin’, in minutes from 8 pm); time of initial Tb recovery (‘Hf’ in minutes from 8 pm) when Tb is maximal after Tmin; heterothermy duration (‘HTD’ (min)= Hf - Hi); and speed of Tb drop and Tb rise (‘V_drop_’ (°C.min^-1^) = (Tb1-Tb2)/(Hmin-Hi); ‘V_rise_’ (°C.min^-1^) = (Tb4-Tb3)/(Hmin-Hf)). Analysis was refined by discriminating periods of normal daily feeding (Mondays to Saturdays), periods of double rations (Sundays), and periods when food was not provided (Mondays).

**Figure 1.**
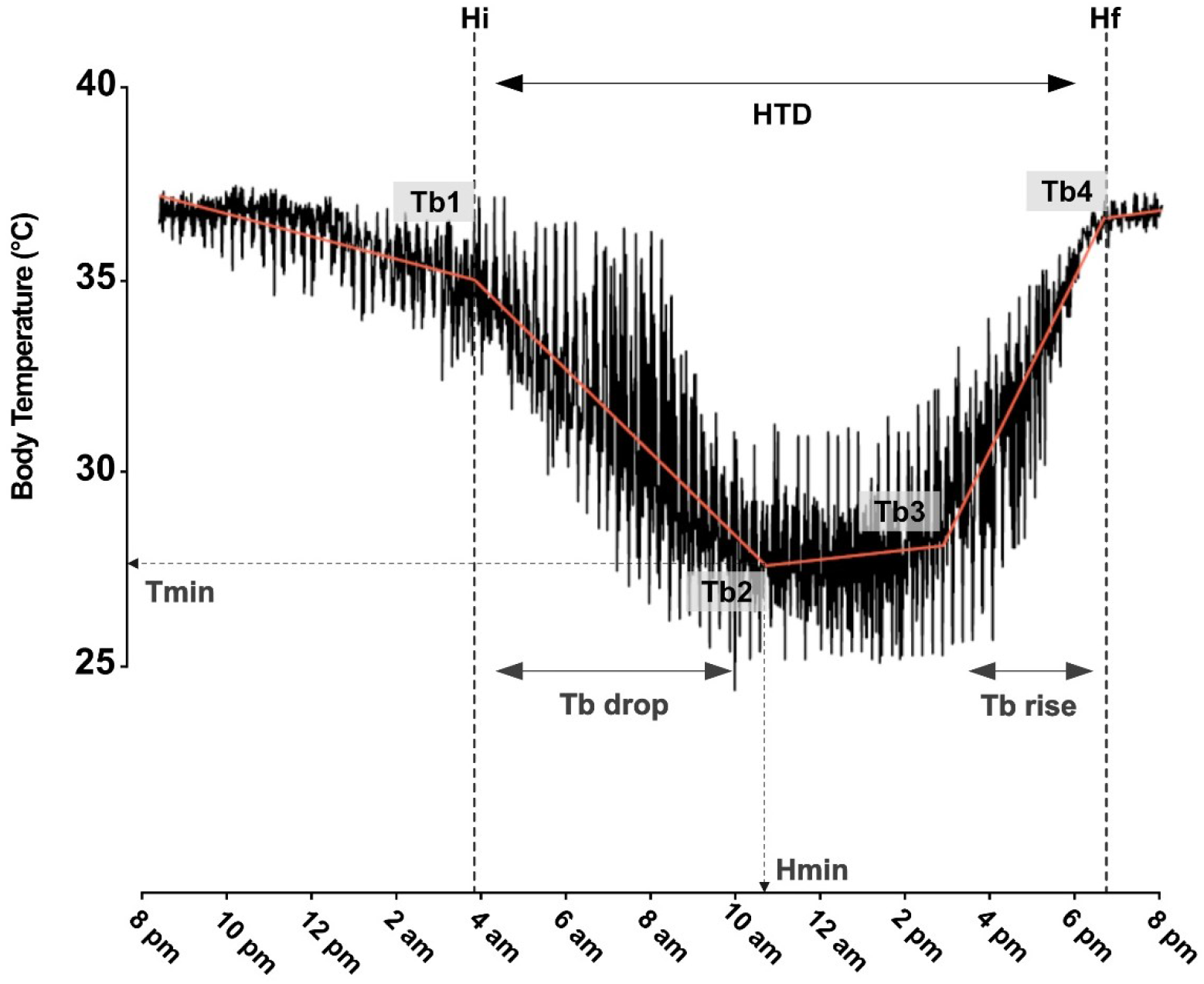
Body temperature (median) over 24h and straight lines (in red) generated through breakpoint analysis. Tb1 and Tb2 are the Tb ordinates framing the linear regression of body temperature fall (Tb drop), with ‘Hi’ being the time of heterothermy initiation. ‘Tmin’ is the minimal ordinate calculated from breakpoints and ‘Hmin’ is the time when Tmin is reached. Tb3 and Tb4 are the Tb ordinates framing body temperature rise (Tb rise) after Tmin, ‘Hf’ being the time of Tb recovering (maximum Tb after Tmin). Each Tb was calculated from the 5 linear regressions as follows: *Tb1 = a0 * Estimate(rank1) + b0 = a1 * Estimate(rank1) + b1 ⇔ b1= (a0-a1) * Estimate(rank1) + b0 (with a being the slopes, and b the intercepts of each regression)*. Heterothermy duration (‘HTD’ in min) was also calculated as the difference between Hf and Hi.

### Reproduction

When the 6 months of SD exposure were over, the animals were switched to summer photoperiod (long days, LD; 14 hours of light, 10 hours of dark, for 26 weeks) and time of estrus was monitored. When females were ready to mate, they were individually put in a cage with two males, one CTL and one CR, taking into account their respective maternal lineages. The lemurs were monitored daily to look for sperm plugs, and females were retrieved from the mating cages 3 days after. Successful gestations were recorded, as well as the number of offspring and their growth rate. Reproductive success was based on the number of pups that survived at least 6 weeks after birth in the case of females. For males this parameter was normalized by the number of mating events in which they engaged (to account for the difference of mating opportunities between individuals). Paternities were assessed from gDNA (extracted from skin samples) using 9 microsatellite markers (see Supplementary materials). Sex ratio of the offspring per adult were calculated dividing the number of male juveniles with the total number of pups.

### Statistical analyses

Results shown are given as median ± standard deviation (s.d.) in Tables. Statistical analysis was conducted with R software v 4.0.0 (R-Core-Team, 2020), and tests were considered significant when p-values were below 0.05. Phenological analysis was performed using GAMM models (‘mgcv’ 1.8-31 package; Woods 2017), when justified with the apparent nonlinear evolution of the explained variable over time, and the statistical significance of the smoothed terms. We averaged each explained variable (BM, Tmin, %Torpor) over the week for the GAMM analyses, as daily parameters were too heavy to compute. Every model presented in Table 1 was confronted to a null model with an ANOVA test, in order to validate that the explained variable did vary with time in a non-linear way, and that it depended on sex and caloric treatment (i.e. that the interaction of the smoothed term with experimental groups is significant; ‘Group’ effect as ‘CF’: CTL females, ‘RF’: 40%CR females, ‘CM’: CTL males, ‘RM’: 40%CR males). An inter-individual effect was integrated as a random variable. Tb variations over 24 hours of the 3 winter periods were also modelized with GAMM to look for the significance of the day of the week effect on the Tb daily profile, in interaction with the experimental group (Table 2).

**Table 1.**
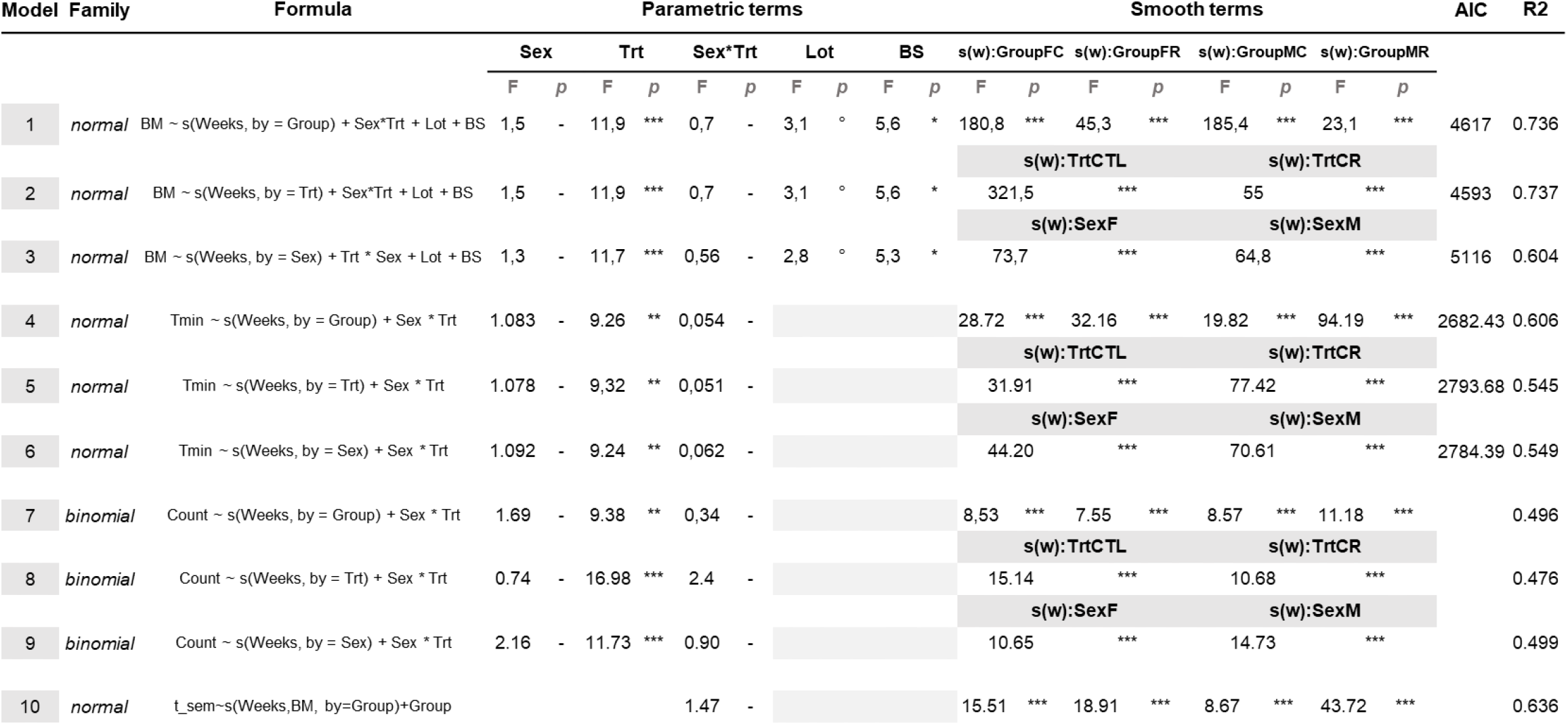
GAMM models of body mass (BM) and daily minimal body temperatures (Tmin) averaged over week, as a non-linear function of time (s(weeks)). The effect of sex and caloric treatment (Trt), which are included in the smooth term as an interaction with time, are investigated by model comparison (AIC coefficient and R^2^). Sex, Treatment, and their interaction (Sex : Trt), are also adjusted as fixed effects to control for differences in intercept. Body size (BS) and the group from which animals originated in the general breeding colony (Lot) are also added as fixed variables when significant. Inter-individual random effect is added in each model. ‘°’<0.1, ‘*’<0.05, ‘**’<0.01, ‘***’<0.001.

**Table 2.**
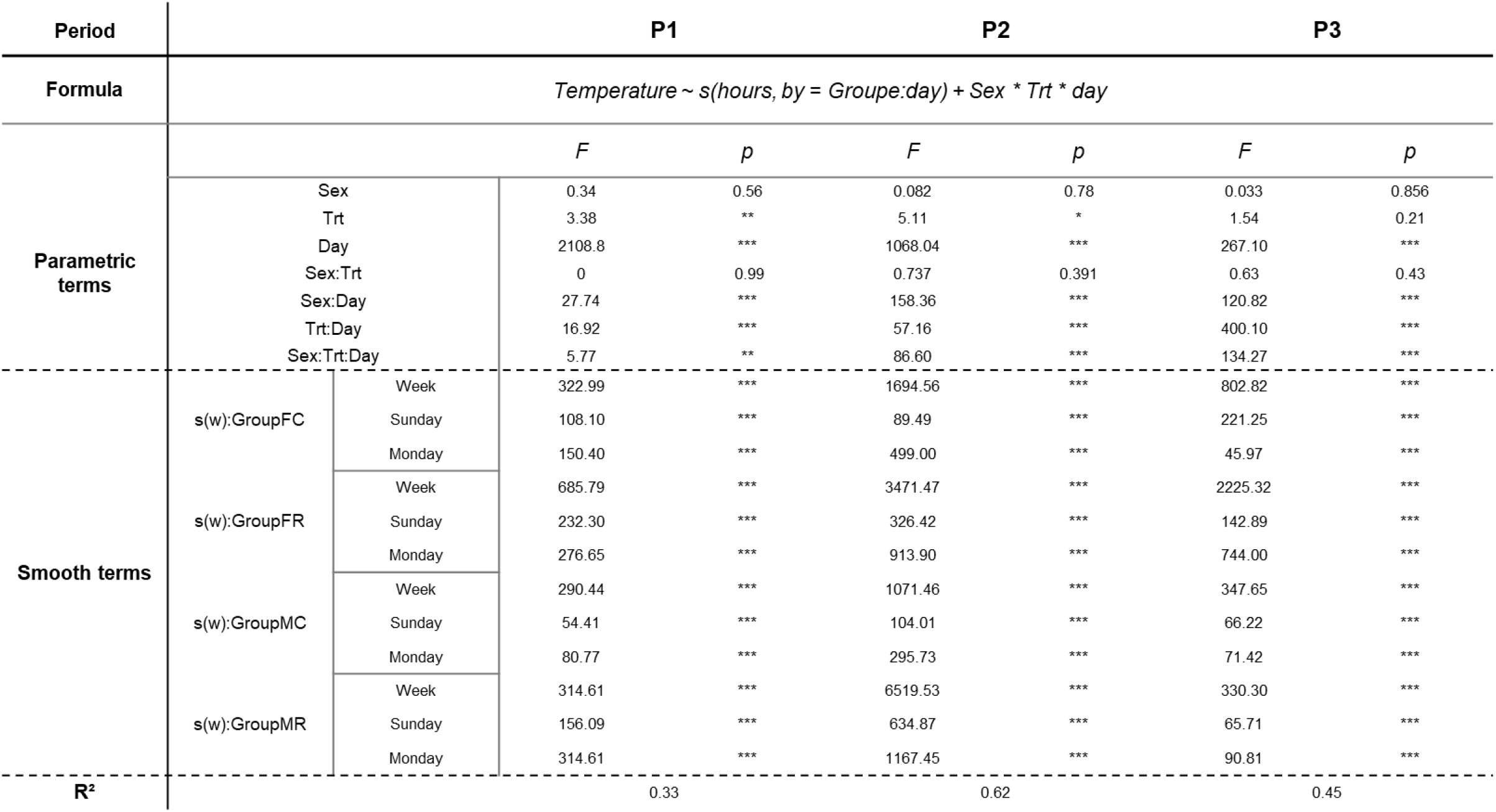
GAMM analysis of Tb over 24h depending on food provision.

For Hmin over winter and quantitative “heterothermic parameters” (i.e. parameters representative of heterothermic profiles over 24 hours, as Hmin, Hi, HTD, Vdrop and Vrise), we applied linear mixed models with random individual effect (see Table 2) to test the effect of sex, caloric restriction, winter time (whether linear for Hmin or as a qualitative factor, i.e. early, mid or late winter), and their interaction. Body mass was included in models only when its effect on the explained variable was significant. For non-Gaussian parameters, generalized mixed models with random effects were used when supported by the data sets. Non-parametric Wilcoxon tests were used for post-hoc analysis.

## 3. Results

### Daily Minimal body temperature vary throughout winter depending on sex and caloric restriction

Daily minimal Tb (Tmin) evolved throughout winter in a nonlinear way (see Figures 2A and 2B), showing significant effects of experimental groups to explain Tmin variations throughout winter, as supported by GAMM models (see Table 1; all groups model explained data better than sex or treatment; model 4: AIC=2682.43, r-sq.= 0.606; only caloric treatment model 5: AIC= 2793.68, r-sq.=0.545; only sex model 6: AIC=2784.39, r-sq.= 0.549). Plotted medians and standard deviations, as well as GMM fitted values, of Tmin over time (in days) showed that females’ Tmin dropped drastically down to 26°C between day 50 to 75, regardless of the CR. In contrast, Tmin levels in males were strongly dependent on food restriction, as CTL males maintained their Tmin around 32°C on the same period, while CR males’ Tmin averaged 25°C. Females differed by caloric treatment after day 110, as CTL females’ Tmin plateaued around 34°C, while CR females Tmin oscillated between 25 and 30°C until the end of winter. Around day 110, males’ Tmin levels were no longer different according to food restriction, increasing toward 35°C to attain a plateau until the end of winter.

**Figure 2.**
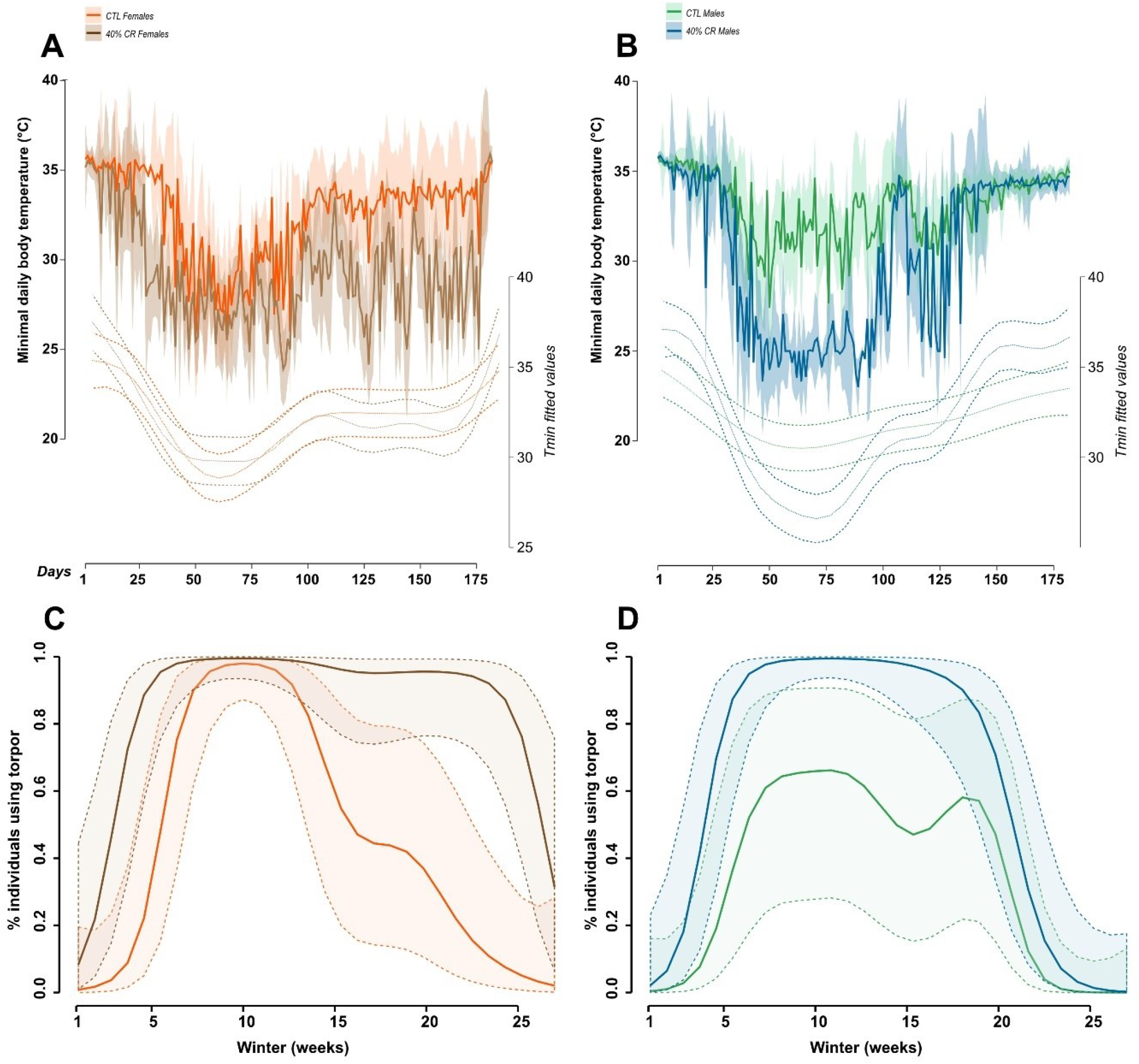
Winter fluctuations of daily minimal Tb (Tmin) (A, B; median +sd in solid lines; fitted values in dashed lines), and proportion of individuals under 33°C of body temperature evolution during winter (C, D; fitted values with confidence intervals from GAMM analysis). A and C: females, where CTL females are in orange and CR females in brown; B and D: males, CTL males are in green and CR males in blue.

Phenological analysis of the proportion of individuals (‘Count’) falling into deep torpor (i.e. for Tmin<33°C) showed strong differences between sexes and caloric treatment (Figure 2C and 2D; models 7, 8 and 9 Table 1). All experimental groups rapidly reached 100 % of individuals using deep torpor around week 5 of winter except for CTL males (Figure 2D), where only a maximum of 60% of the animals used deep torpor during the experiment. In females (Figure 2C), CTL and CR groups differed on the duration of the maximum plateau, as less and less CTL females entered deep torpor shortly after mid-winter (∼week 13), while all CR females used deep torpor until the end (∼week 25). CR males on the other hand ceased to use deep torpor sooner around week 20 (7 weeks before summer transition), as did CTL males.

While BM did not improve GAMM models of Tmin over time as a linear quantitative variable, its integration in interaction with time in the smooth term of the model significantly explained Tmin variations (Figure 3, model 10 in Table 1). This allowed to visualize that BM had not the same impact on Tmin over time depending on the sex and caloric treatment. CTL Females and males had very different profiles, as bigger females (BM>100g) entered deeper torpor especially during early to mid-winter (Figure 3A), while males had a relatively constant Tmin, regardless of both BM and winter time (Figure 3C). Interestingly, the highest Tmin values (Tmin>34°C) were expressed by both very heavy (BM> 140g) or leaner (BM<100g) females during the second half of winter. CR individuals had a flexible Tmin depending on BM and winter progression. CR females weighing 80 to 100g entered into deeper heterothermy (Tmin<28°C) than bigger ones, especially between weeks 5 to 25 (Figure 3B). However, CR females weighing less than 80g were those exhibiting the highest Tmin values (Tmin>34°C) during the second half of winter. In CR males, individuals under 160g experienced bouts of deep torpor under 28°C between week 5 to 20, but warmed up rapidly (Tmin>30°C) at the end of winter if they were under 120g. However very heavy animals (over 150g) were observed only in CTL groups, and these fitted values were unlikely to be observable in CR groups.

**Figure 3.**
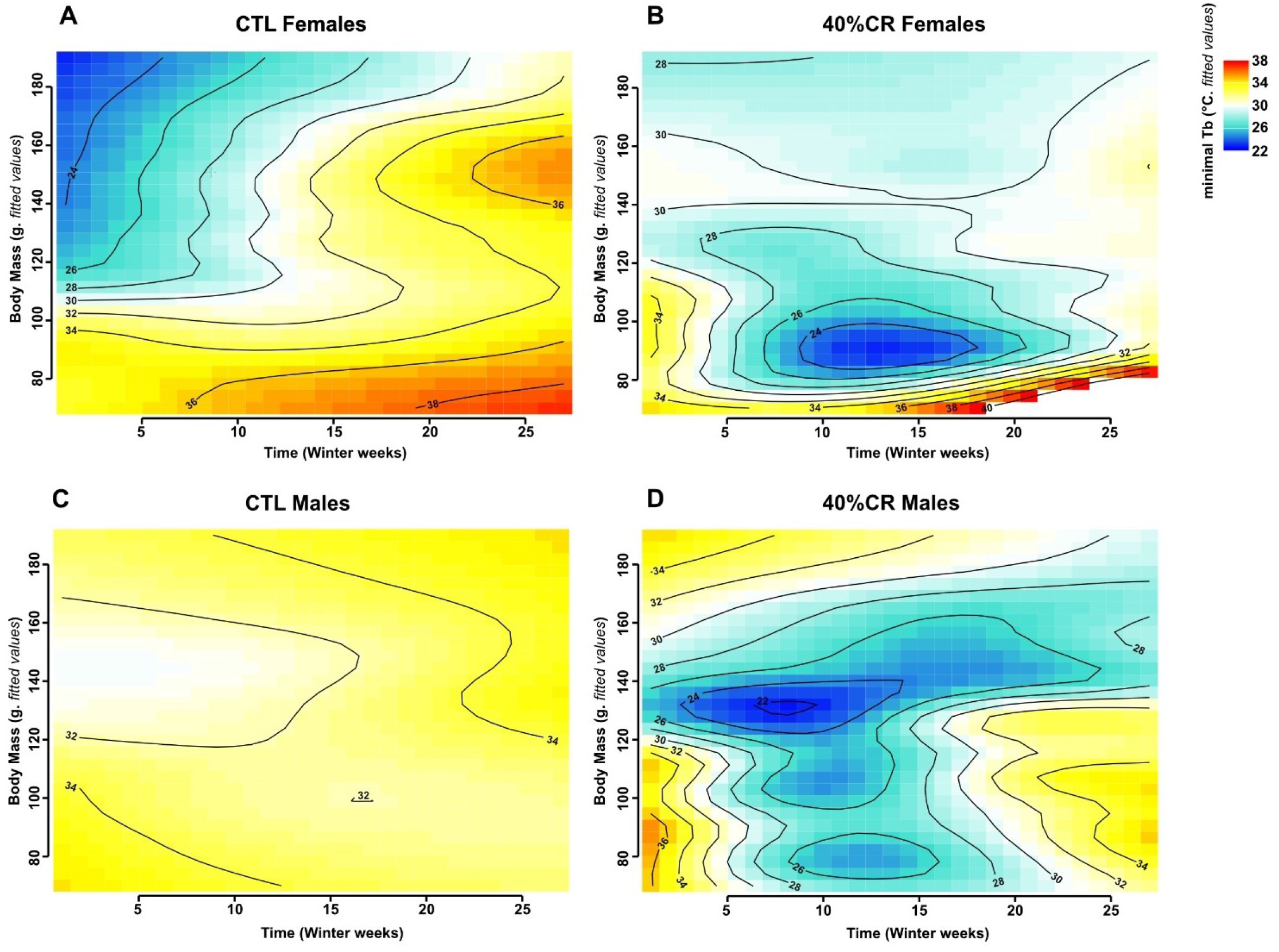
Minimal Body Temperature variations (fitted values after GAMM modelization, Table 1, model 10) as a function of time (in weeks) and Body Mass. Tmin levels are represented by color graduations from dark blue (22°C) to red (38°C). A. CTL females; B. CR Females; C. CTL males; D. CR males.

### Daily Tb profile depends on sex and feeding schedule

GAMM analysis of Hmin over the 27 weeks of winter did not reveal a significant contribution of s(days) smooth terms. Overall, Hmin variation over winter was linear, and Hmin was quite stable throughout winter in every group and occurred between 8am and 9am (Figure 4A and 4B). Time (in days), BM, and Tmin had a significant effect on Hmin, as it slightly decreased over time (meaning Tmin was expressed earlier at the end of winter), increased with BM (the fatter the animals, the later Tmin occurred), and decreased with Tmin (the deepest the torpor, the later Tmin occurred).

**Figure 4.**
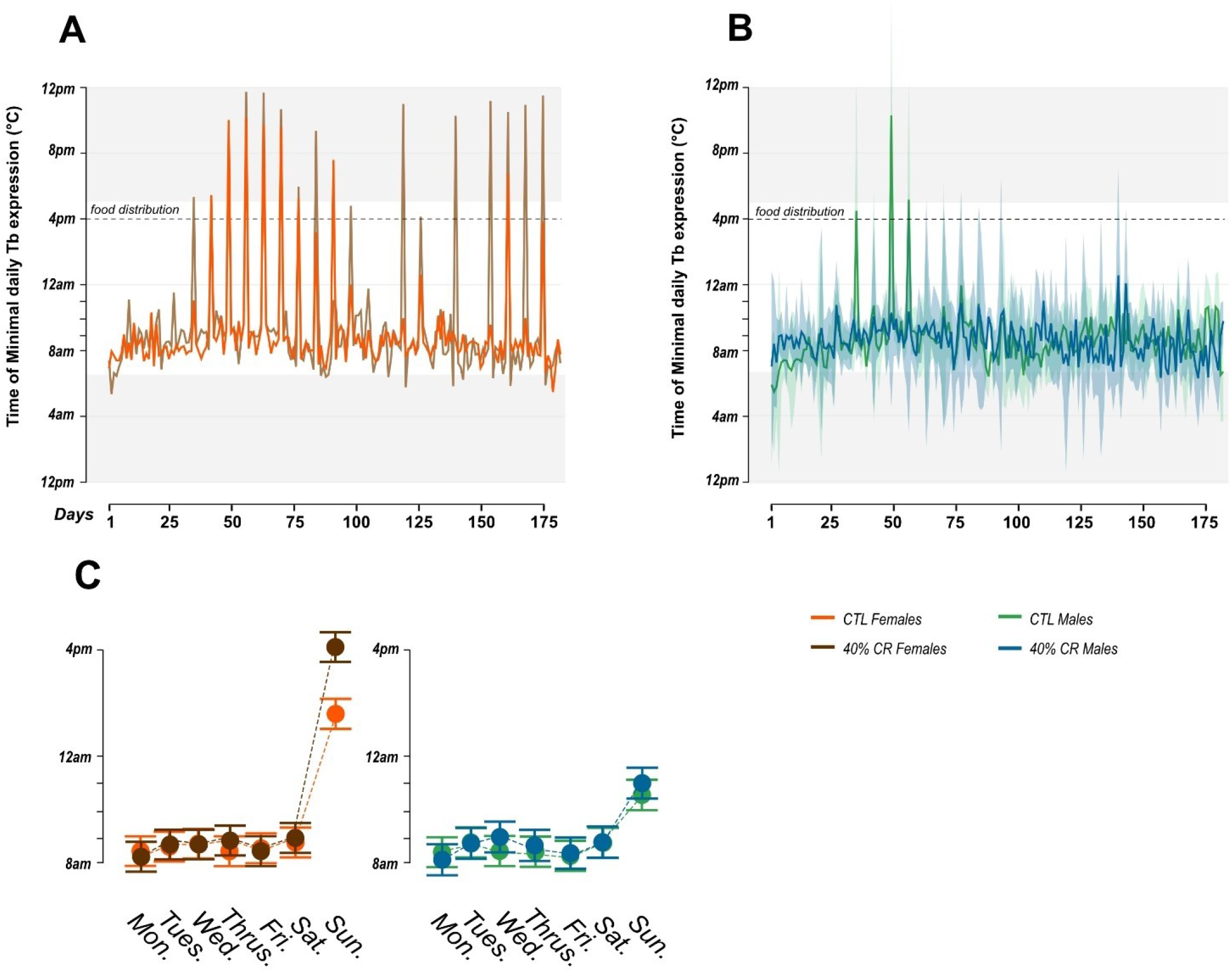
A, B: Time of daily minimal Tb expression (Hmin) over winter (median + sd) and C: graphic representation of time of daily minimal Tb expression (Hmin) modelization by days of the week (lmer): *Hmin∼ time (in days) + BM + Tmin + Day of the week*Group + Lot + (1*|*Individual)*.

However, day-by-day analysis showed that Tmin expression was drastically advanced in a cyclic way (Figure 4A), especially in female groups. This phenomenon happened every Sunday, as animals were fed twice as much on Saturdays for the weekend, leaving Sunday without new food supply (see Methods). The statistical analysis confirmed that females expressed their minimal Tb (Hmin) significantly earlier on Sundays, while males did it at a much lesser extent (Figure 4C; Day:Sex: Chisq= 128.40, p<0.0001).

Further analysis was then performed on daily Tb profiles (Heterothermic profiles, ‘HP’), depending on food distribution (either simple, double or no food ration, whatever their caloric treatment). GAMM analysis showed high significance of food distribution on daily Tb profiles, depending on the experimental group, for all the three periods of winter (see Table 2, Figure 5). Overall, intercepts were significantly lower when no food was distributed on Sundays (Figure 5C, 5F and 5I), meaning that Tb decreased earlier compared to the other days of the week. Smoothed curves showed that torpors were also more profound during these days, with Tb reaching lower values. Models smoothed by experimental groups (Group: Day) (Figure 5) had a lower AIC and higher R^2^ than models smoothed by only sex or caloric treatment, for all three periods (Table S2). This emphasizes the fact that each sex expressed a specific torpor profile, not only depending on the caloric treatment, but also depending on the ‘unexpected’ acute modification of the ration during the week, i.e. double ration or no food distribution. During early winter, food distribution seemed to have little effect on discriminating Tb profiles of each experimental group (Figure 5A, B and C). During mid-winter, CR males underwent deeper torpor (as already shown with the Tmin analysis), and this effect was preserved whether a double ration or no food distribution was given, as all groups modulated the intensity of their torpor bouts in the same direction (lower Tb while no food distribution, higher Tb with double ration). When food was not distributed however, females seemed to enter heterothermy sooner than males, at a slower pace (figure 5F). During the late winter period (Figure 5G, H and I), food distribution modalities allowed to discriminate females from males, as females exhibited lower Tb than males, especially with simple ration or no food distribution. Also, the less food distributed (no food or simple ration), the earlier CR females expressed heterothermy, with few modifications for the other experimental groups, which strengthens the idea that females’ thermic response is more flexible than males’ when it comes to food availability, especially at the end of winter.

**Figure 5.**
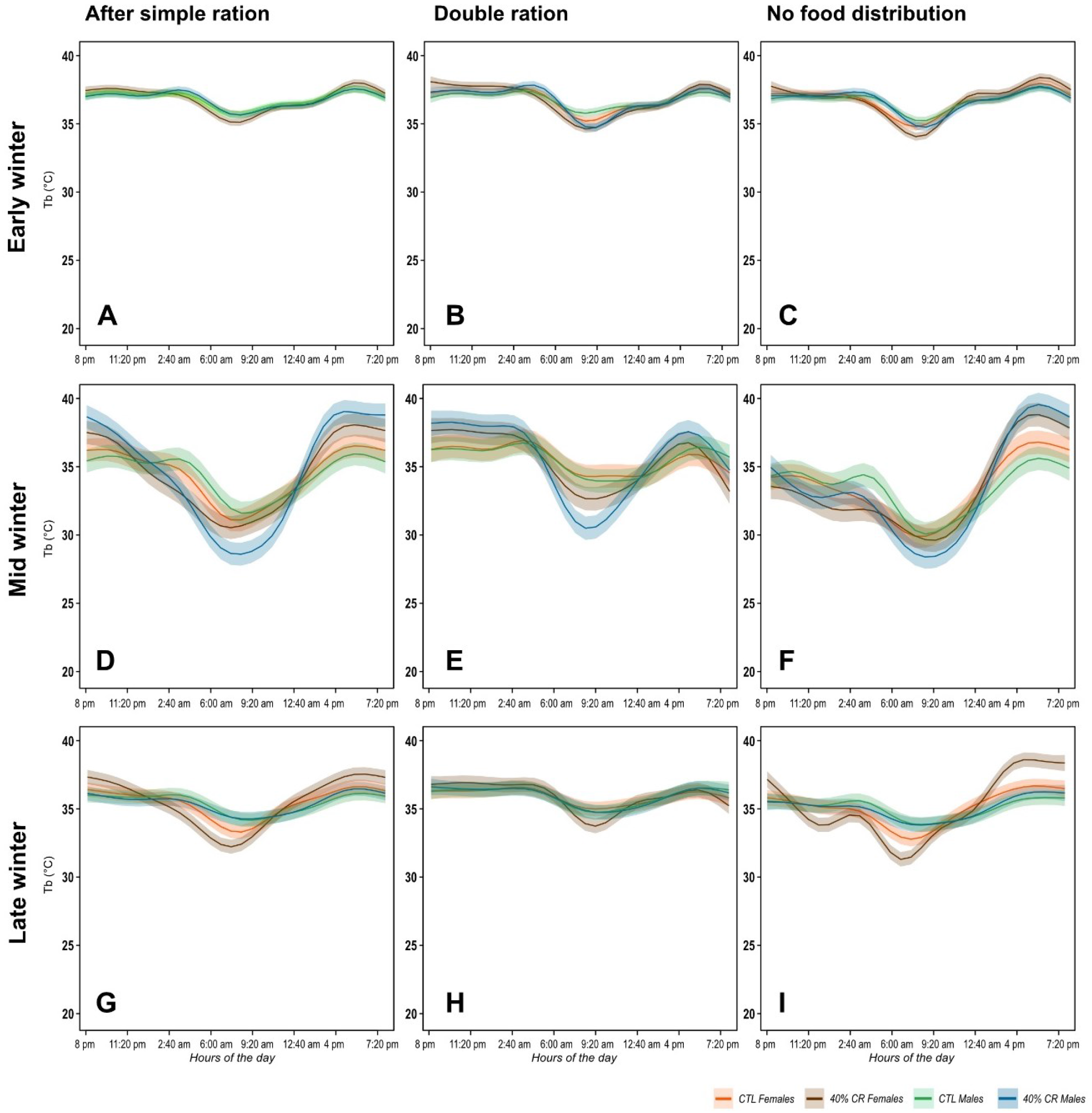
Tb over 24h depending on the period of winter. (A, B, C: Early winter; D, E, F: Mid-winter; G, H, I: Late winter) **and food distribution** (A, D, G: simple ration; B, E, H: double ration; C, F, I: no additional food). Orange: CTL females; Brown: CR females; Green: CTL males; Blue: CR males.

To better define which modality of Tb profiles differed depending on sex and caloric treatment, heterothermic profiles were extracted for all individuals depending on the 3 periods of winter. During the week (Monday to Friday), analysis of daily Tb variations over the 24h cycle showed that minimal Tb (Tmin) dropped between the beginning and the middle of winter, according to the feeding regimen and regardless of the sex (which confirms the analysis of Tmin variation over winter; Figure 2A and B), as CR animals showed much lower Tmin than CTL animals during mid-season (CTL Tmin= 31.71 ± 2.2 °C, CR Tmin= 26.71 ± 2.28 °C, W=134, p-value<0.0001). However, Tmin increased during the second half of winter at the same extent in males regardless of the CR, while females Tmin variations still depended on food restriction (it increased in CTL and remained low in CR females; CTL Females= 34.08 ± 2.84 °C, CR Females= 30.39 ± 2.33 °C, W= 31, p-value=0.04; CTL Males=34.4 ± 0.59 °C, CR Males= 34.58 ± 0.85 °C, W=72, p-value=1). Thus, at the end of winter females seemed to maintain flexibility in the regulation of their body temperatures in response to CR, while males showed unconditional recovery of a normothermic profile, whatever their food regimen. Modelization confirmed that each sex had a specific response to caloric restriction depending on the period of winter (Period:Sex:Trt Chisq= 20.8, p-value<0.0001). Duration of heterothermy (HTD) varied with the period of winter and with CR, but sex did not have a significant effect (Figure 6B; Period:Trt Chisq=9.1, p-value< 0.05). Speed of entry in heterothermic state (V_drop_) increased in males from early to mid-winter regardless of the feeding treatment. The response differed in females and according to CR as V_drop_ remained stable in CTL condition but decreased between early and mid-winter in CR females (Figure 6C; Period:Sex Chisq =25.61, p-value <0.001). At the end of winter V_drop_ decreased in both male groups, while it increased in females, discriminating the two sexes regardless of the caloric treatment. Concerning speed of normothermia recovery (V_rise_) however, caloric treatment had an effect on both sexes (Period: Sex:Trt Chisq= 109.3, p-value<0.001). Vrise was relatively stable during the first half of winter, except for CR males whose V_rise_ strongly decreased (animals regained normothermic status less rapidly). It decreased in the second half of winter for CR animals, increased in CTL males and remained stable in CTL females (Figure 6D). Time of heterothermy initiation (Hi) began earlier in the middle of winter compared to early and late winter, especially for CR animals, regardless of the sex (Period:Trt Chisq=13.2, p-value<0.01). Time of normothermic recovery (Hf) was very stable throughout winter for all groups, as animals woke up when fed, before the lights went off (Figure 6F; CR males appeared to recover earlier than the others, but this was not significant).

**Figure 6.**
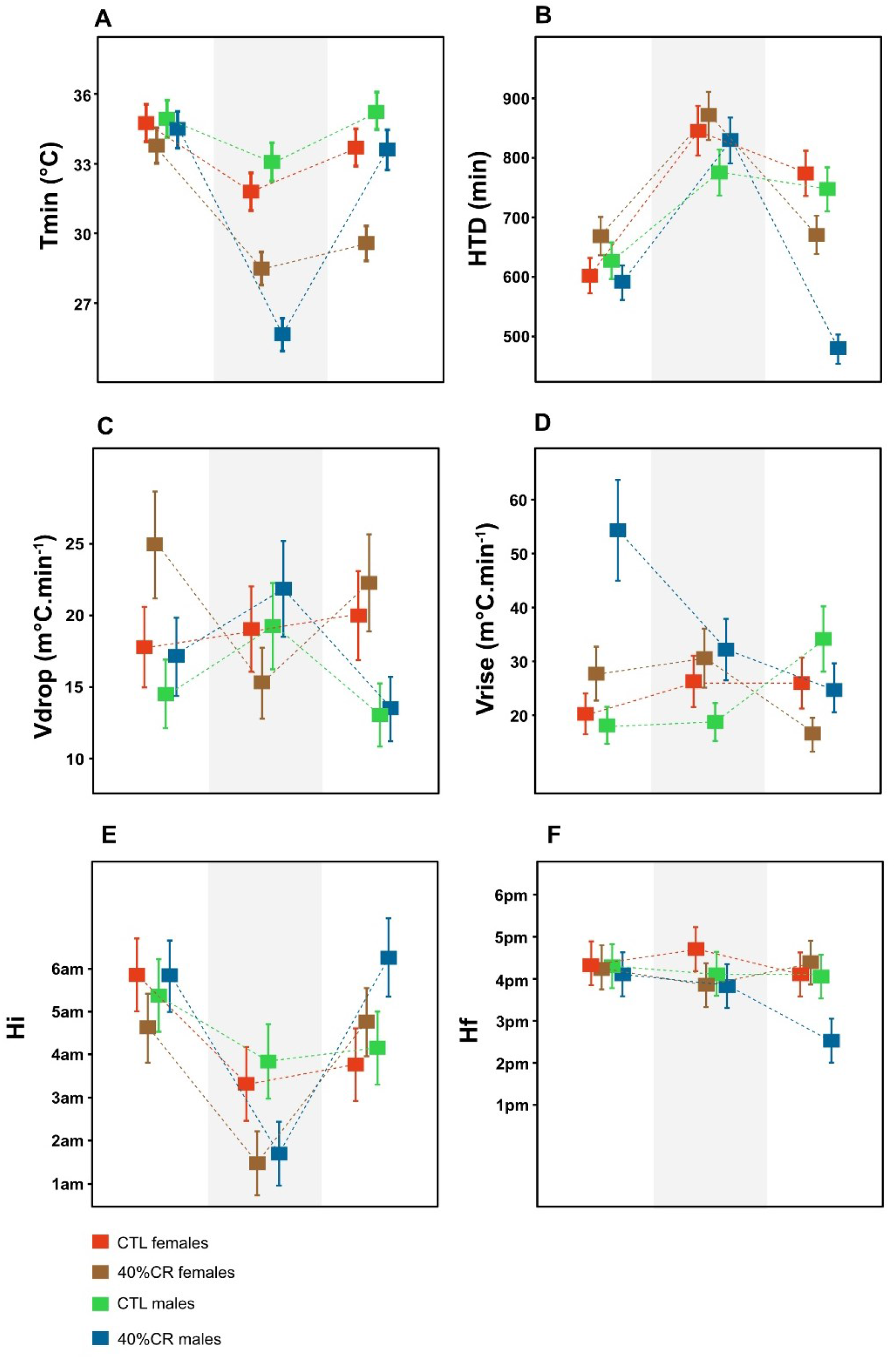
Least Means Squares of different parameters describing heterothermic profiles depending on sex and diet at the beginning, middle and later winter. A: minimal body temperature (Tmin); B: Heterothermy duration (HTD); C: speed of Tb drop before Tmin (Vdrop); D: speed of Tb rise after Tmin (Vrise); E: Time of heterothermic initiation (Hi); F: Time of normothermic recovery (Hf).

### Body condition fluctuations during winter depend on caloric treatment, but not on sex

The relevance of GAMM modelization for the analysis of BM fluctuations was confirmed with the high significance of smoothed time variable, in interaction with experimental groups (Table 1: s(weeks):CF p-value< 0.0001; same result for each group). Animals all consumed their daily food rations entirely. BM fluctuations were identical between sexes throughout winter in function of the caloric treatment (Figure 7A and 7B), as CTL individuals gained weight at a high pace in the beginning of winter to reach a plateau at mid-winter. In contrast, CR animals gained BM more slowly during the first half of winter and lost weight slowly afterwards, independent of sex. However, females had an overall higher BM than males, but this was erased when reported to body size (BS), as females were 1 cm longer than males (Females BS were 13.8 cm vs. 12.8 cm for males; p-value<0.001). Withdrawing sex effect in interaction with the temporal smooth term allowed to discriminate a better model to explain BM fluctuations (see models 1, 2 and 3, Table 1; Group interaction AIC=4617 and r-sq. =0.736; only caloric Treatment interaction AIC= 4593 and r-sq. =0.737; only sex interaction AIC= 5116 and r-sq.= 0.604), which strengthens our interpretation of BM variations being only influenced by time and feeding regimen, regardless of the sex.

**Figure 7.**
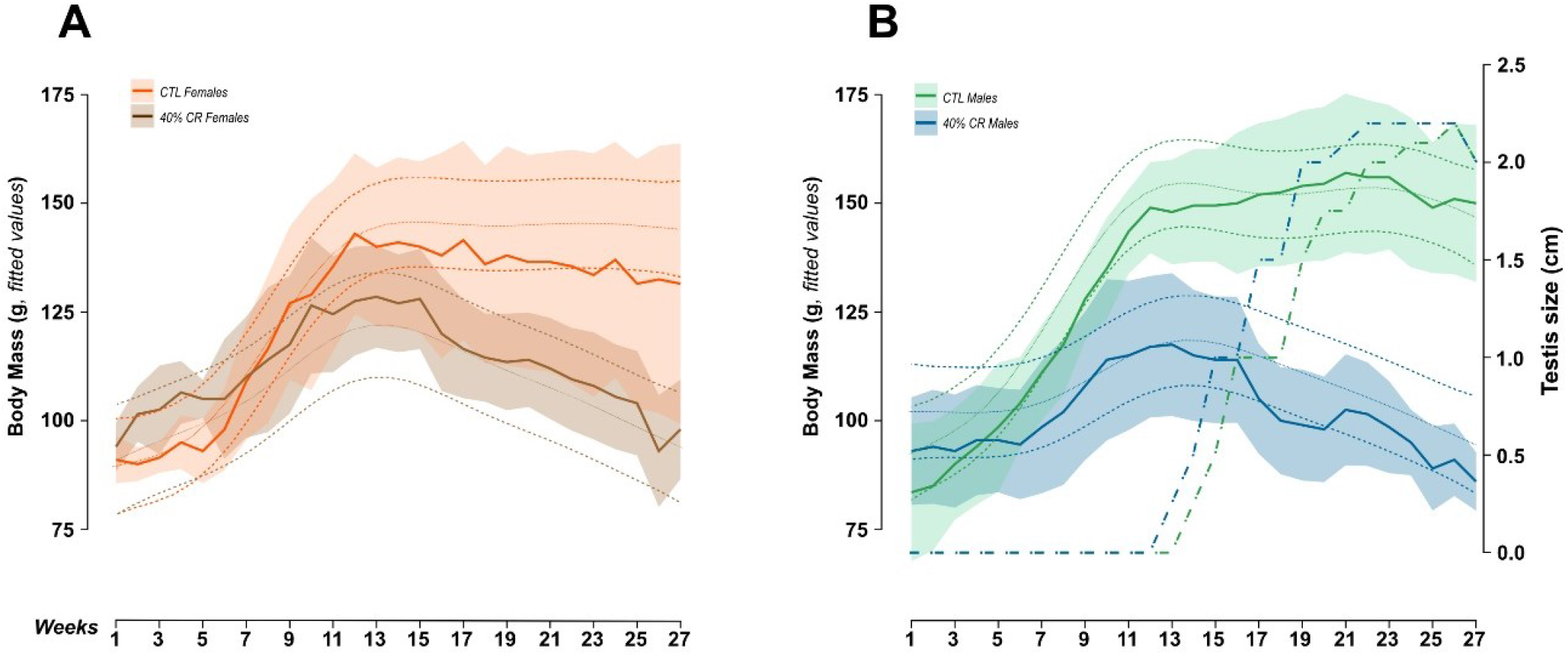
Winter fluctuations of body mass (median +sd in solid lines). A: females, where CTL females are in orange and CR females in brown; B: males, CTL males are in green and CR males in blue. Testis growth over winter is represented in dashed lines.

### Reproductive success

Testis growth was continuous and occurred with a similar rate in CTL and CR males from week 13 until week 23 of winter (Figure 7B). In CR males, the onset of testis growth was congruent with the beginning of BM loss (no BM loss in CTL animals observed in the present data). In females, neither estrus during the entire winter, nor delay in estrus after the transition to LD were observed, depending on the feeding regimen (CTL females= 8.5 ± 4.3 days from photoperiodic change; RC females = 9 ± 4.4 days, W=20, p= 0.81). Six mothers (3 CTL and 3 CR females) produced offspring, with 6 fathers (2 CTL and 4 CR males), and 2 litters had multiple paternities (details of mating groups are provided in supplementary material). The number of offspring produced and which survived did not differ significantly according to sex and CR (CTL females = 1 ± 0.53 and CR females= 1.16 ± 0.66, p= 0.86; CTL males = 0 ± 0.52 and CR males = 0.83 ± 0.60, p= 1). However, total mass of the litters at birth and at 6 weeks tended to differ between mothers depending on the caloric treatment applied during winter (CTL females = 13.6g ± 2.7g at birth and 91.2g ± 5.5g at 6 weeks vs. CR females = 14.5g ± 3.1g then 104.7g ± 6.3g, W=0, p=0.07 and W=0, p=0.1 respectively). Neither mothers, nor fathers conceived litters with a bias in sex ratio, and CR had no significant impact on this parameter (CTL females =0.25 ± 0.5, CR females = 0.5 ± 0.3, W= 5.5, p= 1; CTL males= 0.25 ± 0.4 and CR males = 0.5 ± 0.3, W=18, p=1). Finally, when the offspring were weaned, mothers’ BM did not differ according to CR or reproductive success (with offspring: CTL =90 ± 29.3 g; CR = 95 ± 16.5 g, W=4, p=1; without offspring: CTL= 111 ± 26.0 g; CR= 128 ± 16.5 g, W=4, p=1; reproductive success vs. failure: W=10, p=0.24).

## Discussion

Because a decrease in body temperature (Tb) follows a decrease in metabolic activity (Geiser et al. 2014), heterothermy is seen as an energy saving mechanism in a context of food depletion. Caloric restriction is known to induce torpor or hibernation in many species, including *Microcebus murinus* (Vuarin et al. 2015; Giroud et al. 2008). However, the intersexual variability of these responses is poorly studied. With this winter follow-up, we assessed sex specific variations of body temperatures in a daily and seasonal basis. Moreover, we aimed to test the Thrifty Female Hypothesis (TFH) in *Microcebus murinus*, which would confer to females greater resistance capacities to food shortage.

### Income males, capital females: thermic strategies in response to CR during winter match sex-specific adaptation to seasonal reproduction

Heterothermy is a complex phenomenon, during which one animal can exhibit strong variations of Tb depending on the tissue (Hirshfeld et O’Farrell 1976; Hetem et al. 2016). This can lead to observations of animals performing demanding activities (such as flight) while experiencing low core temperature (Canale, Perret, and Henry 2012; Hirshfeld and O’Farrell 1976; Willis, Brigham, and Geiser 2006). Geiser and collaborators highlighted the necessity to distinguish torpor mechanisms from pathologic hypothermia, and provided tools to recognize active control of metabolic rates and the associated levels of body temperatures (Geiser et al. 2014). In our study, housing conditions, and more specifically ambient temperature (maintained constant at 24-26°C), prevented the animals from suffering of uncontrolled Tb variation, showing rapid autonomic recovery to the normothermic state, unlike passive warming up in case of pathologic hypothermia (Geiser et al. 2014). However, these favourable conditions probably prevented us from measuring deep torpor profiles, and might have contributed to buffer the differences between the experimental groups. Indeed, the minimum levels of Tmin recorded in our study plateaued at a minimum level of 24°C, as expected. Nevertheless, we witnessed many different profiles of daily Tb variations in our long-term study on wintering captive *Microcebus murinus*. Therefore, we combined two approaches to describe these heterothermic profiles, first by working on a continuous scale of Tb, to acknowledge nuances in thermic responses throughout winter other than “above” or “under” a determined threshold, as discussed by Hetem and colleagues (Hetem et al. 2016). Second, we applied the 33° threshold, which is often used (Canale et al. 2011; Giroud et al. 2008; Vuarin et al. 2015), to distinguish shallow (yet clearly defined by Tb dives) from deep torpor bouts, to further explore the effects of sex and caloric treatment on the pattern of heterothermic profiles.

Males and females displayed different fluctuations of minimal body temperature throughout the short-day (SD) period depending on the caloric treatment (Figure 2A and 2C). During the first half of winter, all females -CTL and CR-showed a progressive decrease in their daily Tmin to approximately 27°C, while only CR males reached these minima (and under). CTL males on the other hand, stayed above 30°C during the entire winter. This was confirmed when we assessed the percentage of individuals exhibiting deep torpor (i.e. Tmin < 33°C), as only 60% of CTL males, as opposed to 100% of CTL females, displayed deep torpor in the first half of winter. CR emphasized the use of torpor in both groups, but did not prevent the arrest of deep torpor occurrence at the end of winter in males. The use of deep torpor thus seems much more mandatory for females, regardless of the feeding regimen, while it remains flexible in males throughout winter. Moreover, the duration of heterothermy (HTD) was lower in males as compared to females, especially in the CTL situation, which strengthens the observation that females used deeper and longer torpor bouts than males, potentially optimizing their energy savings.

Moreover, we observed a high modulation of the thermic response in females when subjected to 24h fasting, on Sundays (Figure 5S). On Saturdays, animals were fed twice as much for the rest of the week-end, but they probably ate all their rations at once, especially for the CR animals. Looking closer at the daily Tb profiles, females entered shallow torpors during the usual times (between 7am and 5pm) on Sundays, following the pattern of weekdays. However, they then entered a second torpor bout, more profound, at the end of the day (beginning at 5pm and continuing until Monday to wake up on that day later at 5pm, when food was provided again). The time of daily Tmin expression (Hmin) appeared significantly advanced especially for CR females (and at a lesser extent in CTL females, and a much lesser extent in males whatever their feeding regimen), meaning that they performed deeper and longer torpors on Mondays until the next feeding event. Males on the other hand recovered normothermia for longer periods than females between Sunday and Monday (no food distribution), and began heterothermia later than females, which may inform on their activity to look for food. Some males still managed to follow the female strategy though, which underlines the high inter-individual variability of torpor profiles, even if it doesn’t mask a sex effect.

Torpor or hibernation, both hypometabolism strategies, have been shown to bring negative side-effects, as the generation of oxidative damage during arousals, or downregulation of the immune system (Luis and Hudson 2006; Bieber et al. 2014; Wei et al. 2018; Landes et al. 2020). As males have the necessity to increase their metabolic rate to promote spermatogenesis at the end of winter (Gagnon et al. 2020; Barnes et al. 1986), it may be more beneficial for them to avoid profound hypometabolism if they can get enough energy from the environment. This is probably the case in our experimental conditions, even for the CR, which still received ∼13 kcal/day, which is much more than in the field at the same period (Thorén et al. 2011). In the meantime, females would be selected to capitalize their energy reserves until summer, no matter the current food availability, especially in an hypervariable habitat like Madagascar (Dewar and Richard 2007). On this topic, “Capital breeders” are opposed to “income breeders” for their ability to rely on energy storage which buffers environmental challenges when high expenses are necessary to achieve reproductive success (Doughty et Shine 1997). Other species showed such sex variability in the use of hypometabolism, as polar bear females which enter gestation only when they are pregnant (Lennox and Goodship 2008). Considering our results and the ones that gave birth to the TFH, where females retain more fat than males during winter, we may refine the capital breeder idea by adding a sex-effect.

### Are female Microcebus murinus thriftier than males?

As formulated in the TFH, the most relevant parameter to determine whether a response is energetically beneficial is body mass. Indeed, deep torpors are triggered to save energy when energy (and water) availability is low. While males and females did not show the same thermic responses to CR during winter, or the same use of torpor under CTL ration, they shared the same energy balance. Even if the amplitudes in BM might slightly differ between males and females, as CTL males got fatter than CTL females, and CR males remained leaner than CR females (effect erased with BS normalization), the BM fluctuations during the whole SD period did not significantly differ by sex, but rather by caloric treatment (Figure 7A and 7B). This goes against the previous observations in the general breeding colony, where males start to lose weight during the second half of the SD 6-month regimen, while females retain their BM until the transition to LD, and even until they start lactating if they were successful in reproduction (Perret and Aujard 2001). The reason for males’ earlier BM loss has been linked to their photorefractoriness to SD inhibitory effect on the reproductive axis and the concomitant recrudescence of testis growth and spermatogenesis occurring around week 13 after the transition to SD (although Perret and Aujard showed a recrudescence around week 16, but this may be due to different measurement methods). In the case of *Microcebus murinus*, male gamete production is likely to be an energy consuming process, as they present all the features describing sperm competition, with a high spermatogenic efficacy and motility and low percentage of defect (Aslam et al. 2002; Harcourt et al. 1981). Moreover, spermatogenesis has been directly linked to shorter torpor bouts, as it demands a higher metabolic rate (Gagnon et al. 2020). This would mean a “double cost” for males in winter, because they first have to spend energy to promote their reproductive success during low food availability, and second must lower their best chance in saving energy (i.e. torpor) in order to perform spermatogenesis, ultimately exhibiting higher BM loss than females.

Here, we rather showed that males not only kept their BM during the second half of winter when submitted to a CTL ration (which was close to an *ad libitum* regimen if considering the high weight animals were able to reach), but also appeared to maintain higher body mass than females by the end of winter (which is not significant however, corrected or not by BS). More, CTL females seemed to lose a bit of weight during the second half of the SD period (in median and standard deviation, but the GAMM analysis plateaued). For CR males however, BM decreased by week 13, at the same time as CR females, which is a surprising result of our study. Indeed, females’ reactivation of the reproductive axis is known to be stimulated by the transition to LD, and they show late oestrus when kept under continuous SD exposure (Perret and Aujard 2001). These features -early SD refractoriness in males’ vs LD stimulatory effect in females “endogenous cycles”-have been shown in different seasonal species (Ball and Ketterson 2008). The central down-regulation of the reproductive axis is stronger in females during SD, but the fact that both sexes share the same decrease in BM during the second half of winter, while females use energy saving strategies until the end of the season, raises many questions: what are the energy expenses of females during the second half of winter that justifies such BM loss compared to more active males? What are the benefits of using deep torpor if not to save fat stores for reproduction?

A first indicator could come from the interaction between Tmin and BM. Indeed, this relationship clearly indicates no influence of BM on the levels of Tmin throughout winter in CTL males, while a positive effect of using deep torpor on body condition was observed in CTL females in early winter (Figure 3A), the heavier females exhibiting the deepest torpor. More, higher levels of Tmin (>34°C) could be displayed by either heavy (>140g) or leaner (<100g) CTL females in late winter, while females with intermediate BM displayed shallower torpor bouts. This suggests that the use of deep torpor in late winter was not as frequent and did not benefit as much as in early winter. Second, a study in wintering males recently showed that activity of brown-adipose tissue (BAT) was enhanced in the second half of winter (Terrien et al 2017). This process, which is mainly relying on lipid utilization, is known to be involved in torpor arousal (Townsend and Tseng 2012). Therefore, stimulation of BAT activity could induce higher costs at arousals, therefore counterbalancing the savings at torpor entrance. Whether this hypothesis is true in females should be investigated in the future.

By the first definition of what should be a thrifty female, i.e. a sex able to retain more fat storage during winter than the other (Jonasson and Willis 2011), *Microcebus murinus* females cannot be filed into this category so far. Moreover, an acute 2-week 60% CR during long days did not show a thriftier phenotype in females after reproductive investment (i.e. at the end of the LD period), but our conclusion could not entirely reject the hypothesis, as it is originally linked to the dry season (Noiret et al. 2020). Nevertheless, the 40% caloric restriction we applied during our winter follow-up is relatively mild compared to the food availability in the wild. The CTL regimen may largely exceed animals’ needs, and the CR diet would just correspond to an adjusted ration, allowing animals to use their fat reserves and avoid too much use of torpor. The “torpor optimization hypothesis” states that it should be used only if fat stores are necessary for survival or reproduction (Humphries, Thomas, and Kramer 2003). In the case of a more intense food shortage, females’ thriftier phenotype could become more visible.

Finally, we cannot rule out the possibility that using hypometabolism in a favourable context (i.e. with food) might be associated with other benefits than only body condition, including soma maintenance. Indeed, we observed lower levels of urinary 8-OHdG, a biochemical marker of oxidative stress, in females than males at the end of winter (unpublished data) but also during summer after the reproductive investment (Noiret et al. 2020). In addition, a recent analysis revealed that reproduction, an energetic process, did not accelerate aging in captive female mouse lemurs as expected, but would even promote longevity (Landes et al. 2019). Altogether, these data could be indicative of a better management of the redox status in female mouse lemurs; whether this feature is linked to the use of torpor should be the topic of future research.

### Sex-specific heterothermic strategies: which impact on fitness?

Although the big differences we observed in males’ BM and thermic strategies depended on the feeding regimen, we found no difference on the time of reproductive axis reactivation and final testis size. In literature, males do not respond as much as females on environmental factors such as food availability at the onset of reproductive activity. In females, oestrus delay was often recorded between different populations depending on food availability and quality (Ball and Ketterson 2008; Hahn et al. 2005), while males’ reactivation were not shifted in time (Caro et al. 2009). In our experiment however, the 40% food restriction was not sufficient to induce significant differences in the time of oestrus after transition to LD, again suggesting a favourable energetic context. However, the onset of the first oestrus was not particularly synchronized between females, and a gap of 2 weeks has been observed between the first and last female to enter oestrus, whatever the feeding regimen. Additionally, CR had no impact on either males or females’ reproductive success, even though CR males had more offspring than CTL males (not significant for the 6 males which became fathers). The only parameter that differed was the total mass of the litter at birth and after 6 weeks, which were higher in CR females as compared to CTL females. The contrary was previously observed in mouse lemurs, where a 40% chronic CR applied on wintering mothers during SD only imposed growth delay in offspring (Canale et al. 2012). It would be interesting to monitor the offspring of our study to verify whether CR has affected the fitness of juveniles. Indeed, a well-known mechanism for phenotype transmission across generations is the epigenetic transmission of physiological traits, in response to environmental constraints (Storey 2015). Maternal adjustment of offspring phenotype, through epigenetic modifications, can further contribute to increase average phenotypic plasticity from one generation to the next (Perez and Lehner 2019) and may have operated in CR mouse lemur offspring.

The 40% chronic CR we applied was definitely sufficient to induce lower BM gain during the first half of winter and higher BM loss during the second half, and sufficient to trigger different thermic response between sexes, but did not impact reproductive success. In consequence, both sexes’ respective response to CR seemed protective enough throughout winter in our captive set-up, which may inform on the adaptive nature of their specific heterothermic strategies.

## Conclusion

This long-term follow-up allowed us to assess sex differences in body temperature modulations throughout winter in response to CR in *Microcebus murinus*, indicative of greater use of deep torpor in females, which however did not necessarily translate into lower BM loss. As CTL males stayed fat and CR females lost BM during the second half of winter, our data does not fully support the TFH, although the hypothesis of other benefits than BM should be explored in the future. Finally, the favourable context of our facility might have buffered the effect of sex and further investigation should be conducted in our captive set-up with additional stressors (i.e. low ambient temperatures) or in wild populations, submitted to much more constraining conditions.

## Author contributions

AN, FA and JT designed the work. AN did the animal work. CK performed the genetic paternity tests. AN realized the analytical experiments and statistical analyses. AN, JT and FA interpreted the different sets of data. AN and JT wrote the manuscript. AN, FA, CK and JT reviewed the final manuscript.

## Funding

This work was supported by the Research Department Adaptation du Vivant of the Museum National d’Histoire Naturelle (MNHN). AN received a Ph.D. Fellowship from the doctoral school of the MNHN (ED227).

## Acknowledgments

The expert animal care and management provided by A. Anzeraey, L. Dezaire, S. Gondor, I. Hiron, and M. Perret is gratefully acknowledged. We also thank Amandine Blin from the MNHN’s UMS 2700 2AD for her expert advice concerning GAMM analyses.

## Notes

### Competing Interest Statement

The authors have declared no competing interest.

