## Supplementary materials for "Why be thrifty? Sex-specific heterothermic patterns in wintering captive *Microcebus murinus* do not translate into differences in energy balance"

***Paternity assessment.***

Template DNA was extracted from tissue samples, using the Qiagen DNeasy Blood and Tissue kit according to the manufacturer's instructions. Tissue samples were portions of ear clippings that were taken for identification of each mouse lemur at (AGE). Extracted DNA was amplified using the Millipore-Sigma GenomePlex WGA kit according to the manufacturer's instructions. Primer sequences for targeting microsatellite loci in *Microcebus* were taken from published sources: Radespiel et al. (2001) Isolation and characterization of microsatellite loci in the grey mouse lemur (Microcebus murinus) and their amplification in the family Cheirogaleidae. Molecular Ecology Notes 1, 16-18; and Hapke et al. (2003) Isolation of new microsatellite markers and application in four species of mouse lemurs (*Microcebus sp*.). Molecular Ecology Notes 3, 205-208. Primers were produced by Integrated DNA Technologies. A florescent label (6-FAM or HEX) was added to the 5-prime end of the Forward primer of every primer pair. Each PCR reaction contained 10ul Promega buffer, 8.2ul water, 2ul each 100mM primer, and 1ul template DNA. PCR reaction was carried out as published for each primer set. PCR product was purified with the GeneJET Purification Kit according to the manufacturer's instructions. 10ul of each purified PCR product was submitted to the Stanford PAN facility for fragment analysis, and was measured using the ROX size standards. Fragment analysis results were imported into Biomatters Geneious for quality control and peak calling. Microsatellite allele lengths of each individual were assigned in Geneious using ROX size standards. At each locus, alleles present in the offspring, the known mother, and the prospective fathers were compared. Prospective fathers not possessing those alleles that were found in the offspring and that could not have been passed on by the mother were eliminated as potential fathers. All individuals were sequenced at the Mm10, Mm39, Mm03, Mm09, and Mm21 loci. For most individuals, these loci provided sufficient information to assign a father. When these first 5 loci did not provide enough information, other loci (C1P3, Mm02, Mm08, Mm22) were sequenced until all but one father could be eliminated.

**Table S1. Microsatellites markers for paternity assessment in *Microcebus murinus*.**

| **Locus** | **Motif** | **Length (bp)** | **Annealing temp** | **Accession number** | **Reference** | **Primer F** | **Primer R** | **Alleles published for M. murinus** |
|---|---|---|---|---|---|---|---|---|
| C1P3 | (TC)26 | 205-261 | 50 | AF280079 | Radespiel et al. (2001) | AGCCGAACACATTTCAGAGG | GTAGTCACACCTGGGCTTGG | 21 |
| Mm02 | (GA)18 | 142-172 | 53 | AF280080 | Radespiel et al. (2001) | TTAACAGGGCCTTCTCCTCAC | AATTGCCCAGTCCACACCT | 10 |
| Mm03 | (GA)18 | 93-149 | 55 | AF280081 | Radespiel et al. (2001) | AGCCTCACTGTTTCAGTTGTGT | GGCAGGAAATGTCATCTGG | 15 |
| Mm08 | (TC)18 | 130-198 | 55 | AF280083 | Radespiel et | CAGTTGGTGAATGGGCTAGG | GAGACCATAATGCTGCAAGTAACC | 29 |

| | | | | | | | | |
|---|---|---|---|---|---|---|---|---|
| | | | | | al. (2001) | | | |
| Mm 09 | (TC)24 | 149-193 | 50 | AF280 085 | Radespiel et al. (2001) | TCTGTCTCATGCC TCTTTGCT | GGGTGTGAAAGACA TTACTCACAG | 18 |
| Mm 10 | (CTTT)3CTT(CTTT)2 CTGT(CTTT)13 | 115-152 | 50 | AF280 084 | Radespiel et al. (2001) | GGGCTCCAATAG AGGCAATAA | CTCCAGCCTAGCCA ACAGAG | 22 |
| Mm 21 | (A)17 | 213–245 | 58 | AY154 669 | Hapke et al. (2003) | TCAATGCATCAAT TAACCACG | CAGTTAACATCCTC AGCAATA | 16 |
| Mm 22 | (CA)16 | 204–240 | 58 | AY154 670 | Hapke et al. (2003) | GATATTTGCAGTG ACGTCAAA | AACTTTGACCCTTC CCAGTA | 16 |
| Mm 39 | (CT)6(AC)16(AT)5 | 153–221 | 58 | AY154 673 | Hapke et al. (2003) | TACACTCTGGGTT ACATAAGA | ATCTTTCATCTTCCT GTCCC | 13 |

**Table S2. AIC and $R^2$ based comparison of Tb over 24h models depending on the smoothed terms.**

| | *s(Group :day)* | *s(Sex :day)* | *s(Trt :day)* |
|---|---|---|---|
| **P1** | AIC 142895.3<br>$R^2$ 0.329 | AIC 143500.2<br>$R^2$ 0.319 | AIC 143742.1<br>$R^2$ 0.316 |
| **P2** | AIC 408344.2<br>$R^2$ 0.621 | AIC 427294.7<br>$R^2$ 0.555 | AIC 413009.8<br>$R^2$ 0.606 |
| **P3** | AIC 217154.4<br>$R^2$ 0.447 | AIC 220148.2<br>$R^2$ 0.423 | AIC 223923.0<br>$R^2$ 0.394 |
